# Murine Radiation-Induced Stomach Pathology from Whole Thoracic Irradiation

**DOI:** 10.1101/2020.03.06.980987

**Authors:** Daniel R. McIlrath, Carlos J. Perez-Torres

**Affiliations:** School of Health Sciences, Purdue University, West Lafayette, Indiana 47907, United States; Purdue University Center for Cancer Research, Purdue University, West Lafayette, Indiana 47907, United States

## Abstract

**Purpose:** Radiation-induced lung injury is a common side effect in the treatment of lung and breast cancers. There is a large focus in the field on leveraging mouse models of radiation-induced lung injury to find novel treatments for the condition. While attempting to irradiate mouse lungs for purposes of creating a radiation-induced pulmonary fibrosis model, noticeable declines in health were observed at much earlier time points than recorded for lung pathology. This was later attributed to stomach pathology observed in CT images and ex vivo dissection.

**Methods:** For this study, we used longitudinal microCT to characterize male C57Bl/6 mice irradiated with a single dose of 20 Gy to the whole thoracic area delivered by an X-Rad cabinet irradiator. CT was performed with respiratory gating at 2 to 4 week timepoints to construct a timeline of pathology leading up to fibrosis and quantify severity of fibrosis afterwards. However, a mouse imaged at the 10 week timepoint showed evidence of stomach distention. These mice were sacrificed and their stomachs removed. Histology was performed on the stomachs using H&E staining.

**Results:** On the CT images, we observed a large, spherical volume of hypointense signal, caudal to the lungs (Figure 1). This correlated with a distended stomach caused by constipation and gas build-up within the stomach. Statistical analysis showed 81% of mice (n=105) died prematurely after irradiation and before significant development of pulmonary fibrosis. Mice sacrificed and dissected showed unpassed bolus as contents of the stomach, and histology showed cell necrosis of the stomach walls.

**Conclusion:** The histology indicated an inability for food to be digested and moved into the small intestine. This lead to a blockage and ensuing stomach distention. Given the severity of the pathology’s consequences, it lead to the mouse’s imminent mortality inhibiting the efficacy of the study. Future studies need to consider careful placement of shields or any beam contouring devices to ensure protection of the stomach given its higher radiosensitivity in contrast to the lungs.

## Introduction

Thoracic irradiation treatments are among the most common in external beam cancer therapies. This is largely due to the high incidence of breast and lung cancer – the two most prevalent cancers in the world. Lung cancer is the most frequently diagnosed cancer with 1.6 million new cases each year^1^ and breast cancer is the second most frequently diagnosed cancer with 1.1 million new cases each year.^2,3^ Proving to be an effective treatment, radiation therapy ideally destroys all of the cancer’s tumor cells while maintaining healthy cells’ integrity. However, the ideals prove to be far from realistic with 35% of thoracic irradiation patients at risk to develop lung injury.^4–6^ Therefore, it is necessary to examine radiation injury to thoracic organs and tissue through pre-clinical models in mice.

To create such a model, accurately contouring the beam becomes vital due to the small nature of the mice’s organs and the nearby relative radiosensitivity of neighboring organs – most notably the stomach. Clinically, this is possible through a combination of multi-leaf collimators and on-board imaging to contour the radiation beam to the desired treatment field with great efficacy. Pre-clinical studies rarely have access to such sophisticated devices and rely on simple radiation machines, such as cabinet irradiators, to deliver the experimental dose. These crude devices, in comparison to LINACs, lack the ability to contour beams as accurately, and therefore, require subjects be shielded to protect areas not being researched.^7–10^

Improper shielding can lead to unexpected off-target effects, particularly of tissues that are radiosensitive. These off-target effects could potentially present a confound that may obfuscate the intended results such as the mortality of the subjects or injury development. Here we report on unintended stomach toxicity in a model of radiation-induced lung injury.

## Methods

*(All animal experiments were approved by the Purdue Animal Care and Use Committee.)*

### Mouse Characteristics

Mice used to develop our model was determined based on previous literature.^7^ Male C57Bl/6 mice were used due to their reported radioresistance compared to female mice and their historically published pro-RIPF characteristics. Radiosensitivity was of concerned because previous trials preformed lead to premature death leading to the decision to use mice that were more likely to survive irradiation. Mice were irradiated at 8 weeks-old and weight was not tracked.

Mice were housed in a facility located on Purdue University’s campus and maintained by Purdue University’s animal care staff. Mice were checked daily by staff and weekly by researcher during imaging dates. Mice that developed severe wounds due to erythema, bullying, or other causes were checked every other day. Euthanasia would be deemed necessary by animal care staff when wounds, after treatment, progressed or remained severe.

### Irradiations

Based on previous studies^7^, mice were irradiated with a single fraction dose of 20 Gy to the whole thoracic region. Irradiations were performed with a 320 kV cabinet irradiator (X-Rad 320, Precision X Ray, North Branford, CT). Mice were sedated with Isoflurane delivered via a vaporization and anesthesia device and then kept sedated via the same device on a five-chambered platform with anesthesia channels (Figure 1[A]). Mice were sedated in groups of five and placed on the platform in the prone position with Gafchromic EBT 3 film (Ashland Advanced Materials, Bridgewater NJ) placed beneath for position validation. Lead shielding was then placed above the mice (Figure 1[B]) with a 15 mm x 15 mm aperture placed above the thorax of each mouse. Each mouse was adjusted individually to optimize positioning for dose delivery. The cabinet irradiator was set to deliver a dose of 20 Gy with a dose rate of approximately 200 cGy/min and a tube setting of 320 kV. Source-to-skin distance was set to be 50 cm and a 0.1 mm Al beam flattening filter was used. Irradiations took approximately 10 minutes to deliver the desired dose given the dose rate. Afterwards, mice were allowed to recover from anesthesia in their cages under observation and returned to the mouse facility.

**Figure 1:**
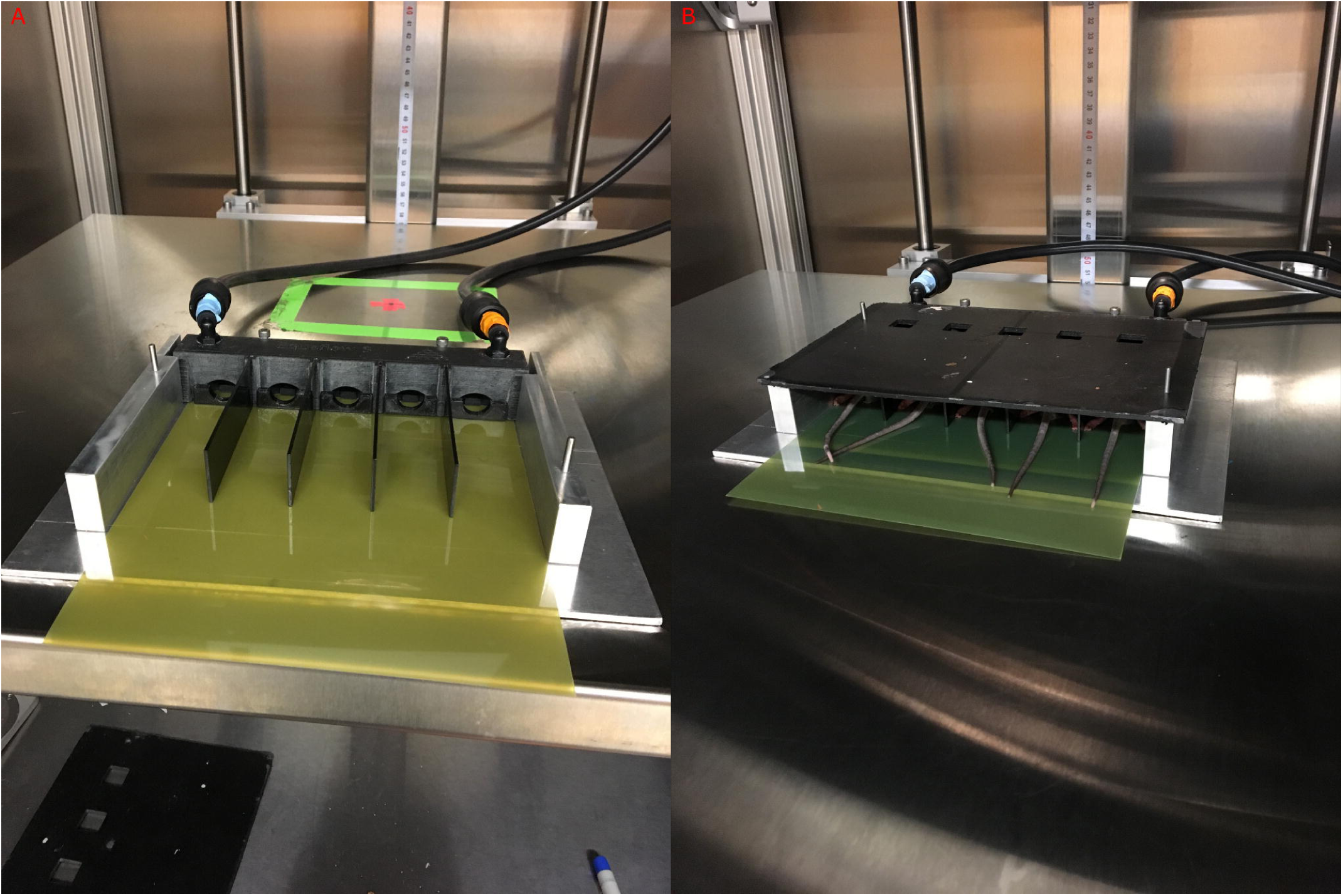
Median Survival Times for male and female C57Bl/6 mice receiving varying Whole Thoracic Irradiaiton doses.

Calibrations for the X-Rad 320 cabinet irradiator were performed by another lab group and dose rate was reported to be approximately 200 cGy/min. Aberrancies in the dose rate were minimal, in reference to the overall delivered dose, and systematic throughout the experiment rendering our results to be unaffected by any deviation in the dose rate.

### Film Analysis

Film irradiated during dose delivery was analyzed using an EPSON Perfection V600 photo scanner and the software program ImageJ. After scanning films, the corresponding files were opened with ImageJ where their RGB channels were separated into three individual images. The green channel offered the most contrast for Gafchromic EBT 3 film, so it was used to validate subject positioning during irradiation.^11^

### Histology

Mice that showed distended stomachs were euthanized using Isoflurane and had their stomachs removed and placed in 4% Paraformaldehyde. Stomachs were then sliced along its longitudinal axis and placed in 70% ethanol. Slices were stained with hematoxylin and eosin by the Purdue Histology Facility.

### New Shields

After noticing this issue, two new shields were created – one that had 7.5 mm x 15 mm sized apertures and another that had 7.5 mm x 7.5 mm.

## Results

### Evidence of irradiation beyond the lungs

The original goal of our study was to evaluate a mouse model of radiation-induced lung injury. As a quality control measure, we utilized radiographic film to image the portions of the mouse that received irradiation. It was noticed that many of the mice were misaligned since the aperture was larger than the size of the lungs, and large portions of liver and stomach were receiving the same dose as the lungs (Figure 2[A]). By changing the size of the aperture within the lead shielding we reduced the amount of non-lung tissue that was irradiated as evidenced by Figure 2[B]. Even so, as the position of the mice were not perfectly consistent between cohorts there might still have been some variation with regards to the localization of the irradiation field.

**Figure 2:**
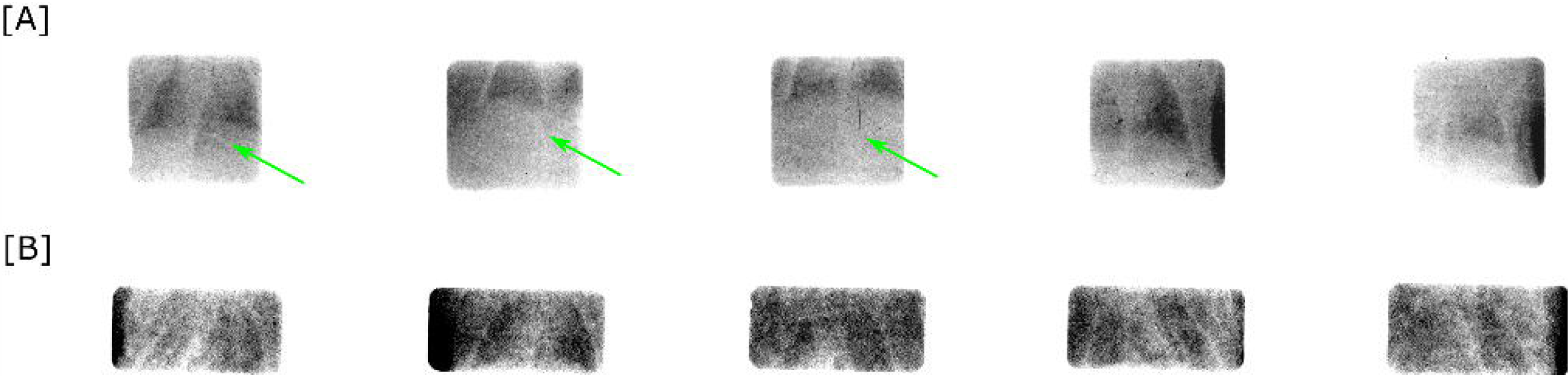
Irradiation setup with [A] a 5-chamber platform and attached to a sedative delivery system with a piece of Gafchromic EBT 3 film slid in between the floor of the platform and the mice. [B] Mice placed in chambers with lead shielding placed above to contour the x-ray beam to irradiate the thorax of each mouse to deliver 20 Gy to only the lungs.

### Mouse Survival

After changing to the new shields, shield 2 resulted in 0% of the mice dying prematurely (n=15) and shield 3 resulted in 20% (n=35) of deaths being premature. These numbers definitely indicated an improvement to our experimental design that allows a greater number of subjects to survive to exhibit the desired pathology. The major consequence of the larger irradiation field was mice died only a few weeks after radiation whereas death due to fibrosis should occur at ∼24 weeks post irradiation at our given radiation dose.^7^ By reducing the radiation field size and therefore reducing the unintended off-target effects, early mortality could be minimized (Figure 3).

**Figure 3:**
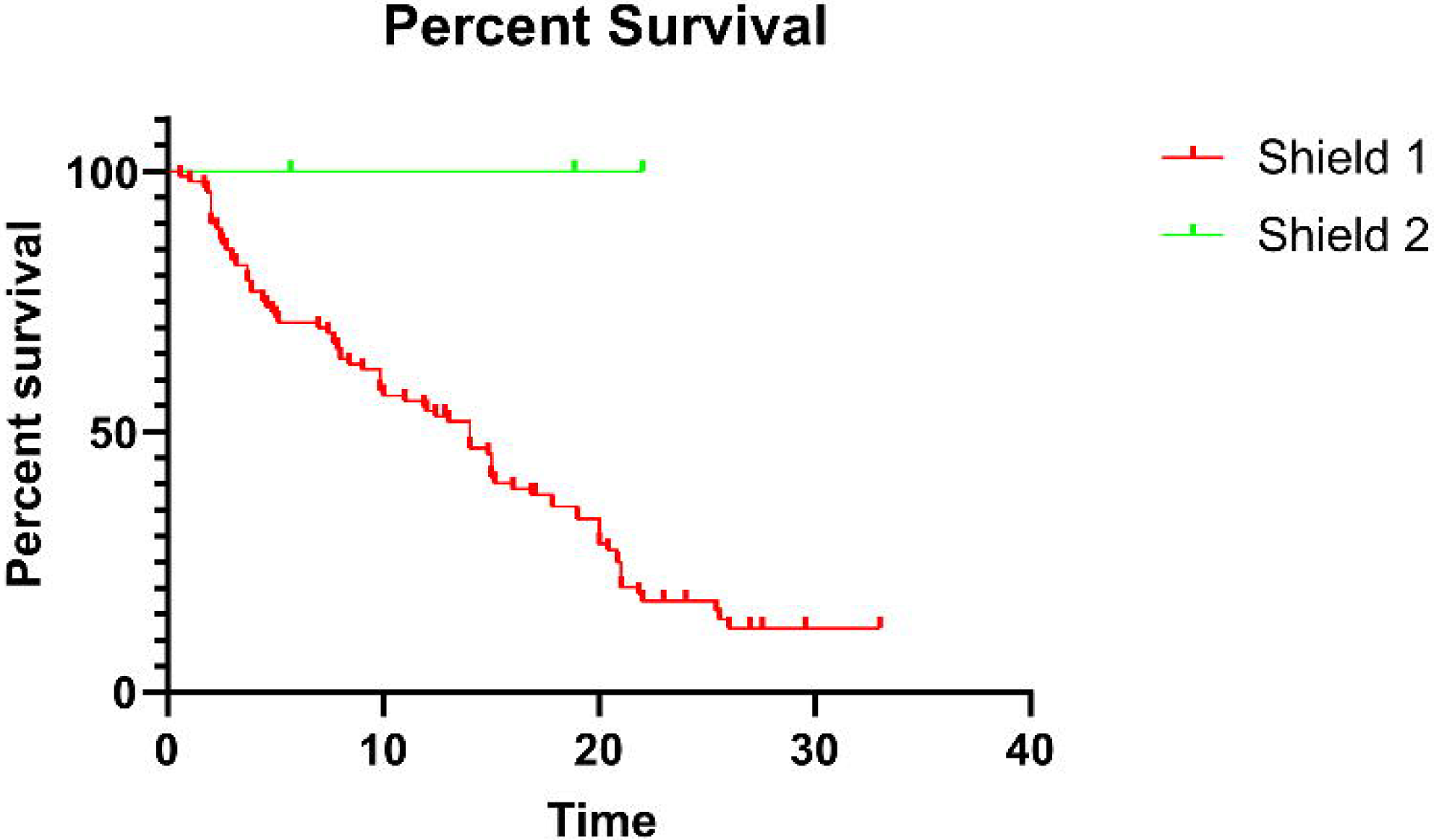
Film of 5 mice’s positions relative to the aperture in the lead shielding. It is evident from these films that positioning of the mice during irradiation was an issue since 3 of the 5 mice had significant portions of their abdomens exposed in the radiation field (green arrows) with [A] shield 1 as opposed to [B] shield 2. This would lead to a large direct dose being received by the stomach resulting in significant injury.

### Stomach Pathology can be identified on CT and verified on histology

The premature deaths in our study are at least partially due to unintended stomach pathology. On CT examination the normal stomach appeared as a minimal hyperdense area with gas pockets or a small hypodense area of air if the dorsal portion is viewed (Figure 4). For a mouse that died prematurely, the stomach is observed as a large hypodense area just caudal to the left lung. This CT hypodensity could be observed as early as 4 weeks post irradiation which is consistent with the timing of premature death. That the stomach is distended and hypodense on CT suggests a large buildup of gas, likely due to the inability to break down food.

**Figure 4:**
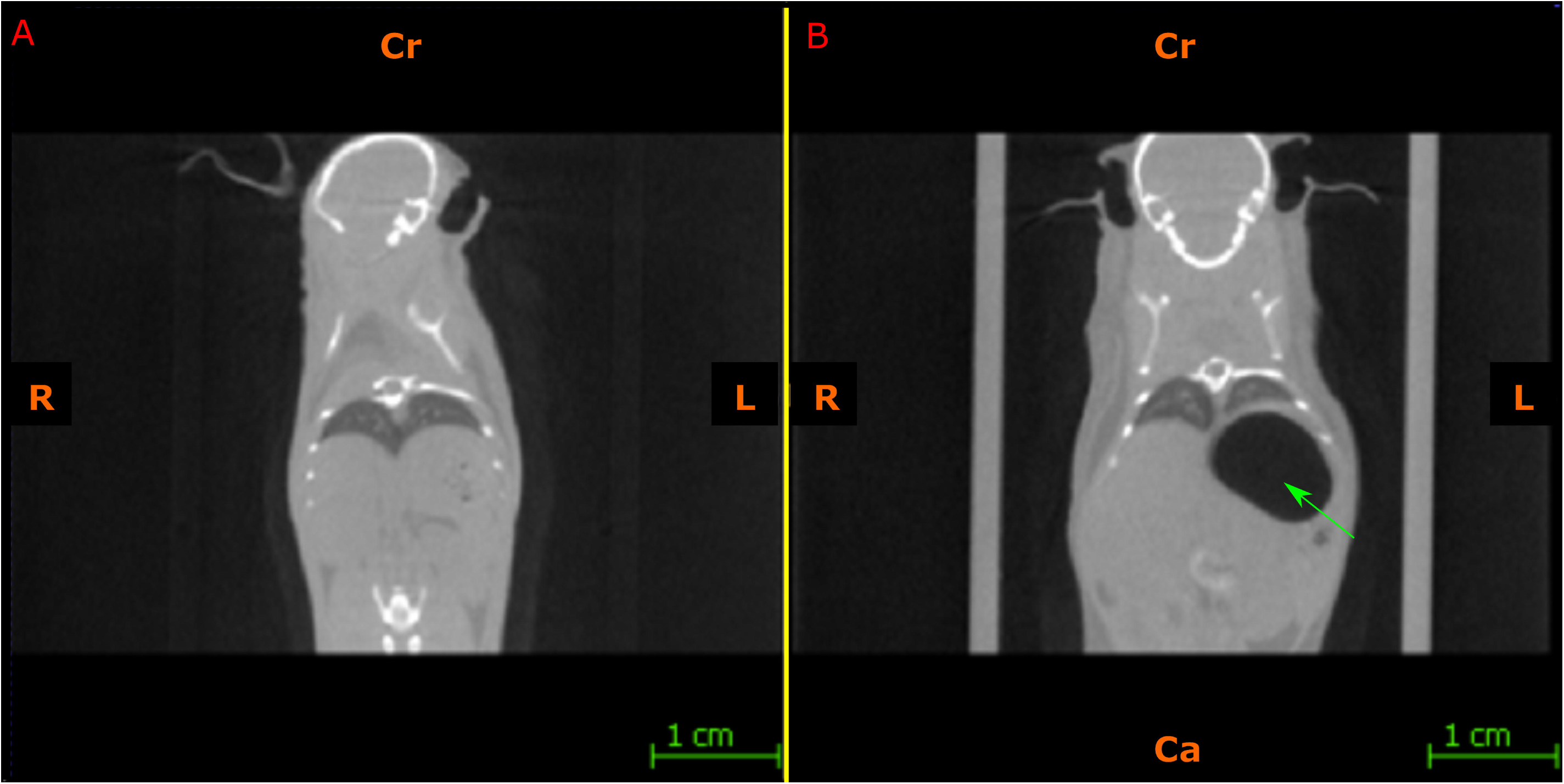
Survival of mice irradiated with shield 1 and 2. Mice survived significantly longer when shield 2 was used in place of shield 1 to irradiate the mice. This led to the conclusion that by minimizing the aperture size, positioning was made easier and a tighter exposure field to the lungs was more possible.

Seen in Figure 5, histology shows major gaps in the middle section of the fundic region of the mouse’s stomach. The basal and superficial layers appear to show little alteration; however, there is structural transformations that appear to extend from the superficial layers through the middle to basal layers. The most radiosensitive cells of the fundic region are the eosinophilic parietal cells of the middle layer which secrete acid for digestion, and the next most sensitive would be the mucosa-secreting cells of the foveolar column of the basal layer.^12,13^ The desquamation of the mucosa most likely lead to ulcerations as seen in Figure 5[B] (green arrow).

**Figure 5:**
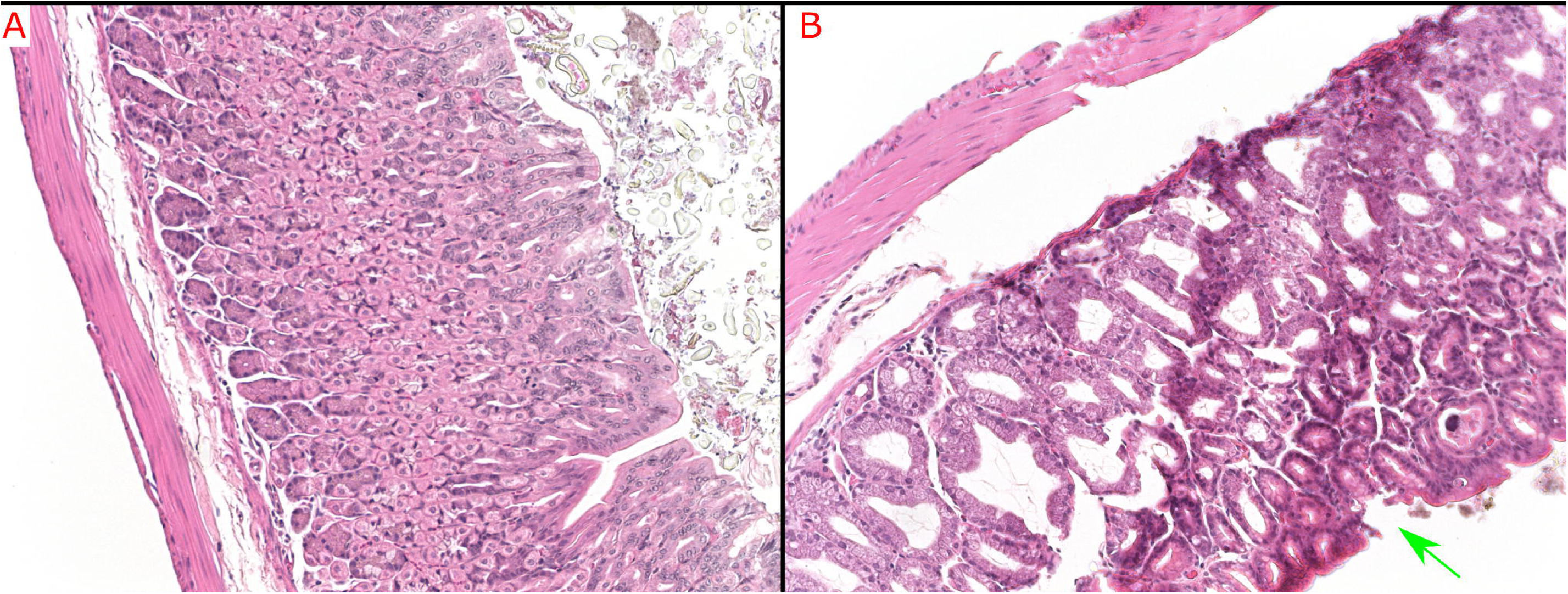
CT image of thorax [A] pre-irradiation and [B] 10 weeks post-irradiation. The mouse’s stomach has a large area of hypodensity caudal to the left lung (green arrow). This area coincides with the stomach and the abnormal size of this area implies the stomach is distended. Although specific dosimetry would need to be done to determine the exact dose received by the stomach, the dose that we delivered led to high enough doses being absorbed by the stomach to cause early injury that jeopardized the efficacy of the study.

## Discussion

Thoracic pre-clinical radiation modeling significantly aids future studies by creating a basis to which experimental factors can be compared. However, these models prove difficult to construct due to the small size of the subject and areas adjacent to the thorax, such as the stomach, easily irradiated unintendedly. The extreme sensitivity of the stomach when compared to the lungs jeopardizes any study that may expose it to the radiation field, thus future research must be mindful of the stomach’s radiosensitivity in order to assure unimpeded results.

From our CT images, it was clear that pathology beyond pulmonary injury was occurring. Stomach distention became salient on CT images of any effected mice and indicated improper radiation delivery. This could be further verified by the films produced during irradiation. When the radiation field size was changed by adjusting the aperture size on the shielding, significant changes in survival were observed. Histology provided even more evidence of the injury to the stomach that aligns with previous studies on stomach radiation toxicity.^8,9^

Due to the salient stomach distention observed on CT images, the stomach was the focus of this tangential observation during lung modeling. Liver was considered as a possible organ at risk but quickly dismissed due to its radioresistance compared to the stomach 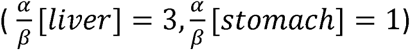^14^ and lack of pathology observed on the CT images.

The gastro-intestinal (GI) tract and other distal structures to the lung were not considered in our study since the misalignment of the mice was minimal enough to not expose extremely caudal features and our misalignment was always exposing the abdomen, so no organs cranial to the thorax were considered.

During external beam treatments, the human stomach often receives residual dose from targeted tumors in the upper gastrointestinal tract or inferior lung. If the stomach receives around 15 Gy in a single fraction, late-effects begin to be of concern despite the volume of the stomach irradiated.^15^ Patients that have received around 67 Gy total through fractionation have reported ulcers occurring post-irradiation.^12^ In a three-fraction stereotactic body radiation treatment (SBRT), dose-volumes are required to be minimized to 22.5 Gy to 5 cc of stomach^15^. In rodents, doses above 14 Gy, delivered in a single fraction to the whole stomach, showed significant risk in the development of acute- and late-effects and eventually mortality^9^. Since doses for pre-clinical lung studies are close to this threshold, it is extremely important to thoroughly protect the stomach during these studies.

## Conclusion

Though we were able to confirm the presence of stomach pathology in a few of our mice that died prematurely, there were also some mice where necropsies could not be performed. Some premature deaths are left out of this analysis since their cause of death was known – overdose of anesthetic during imaging – although this number was small (n=8).

The premature deaths, presumably caused by stomach pathology, were a major challenge to our primary study focused on radiation-induced pulmonary pathology. This challenge was further exacerbated in the case of the mice for whom necropsy was not feasible. Without necropsy it was not possible to ascertain whether death was due to lung pathology or unintended pathology. This also further demonstrates the limits of using survival as a metric for specific form of radiation-induced pathology. For this reason, it is important to create adequate protection and develop a mold that provides more precise subject placement in future lung studies to prevent a loss of potential data and time.

Unfortunately, since this was an unplanned pathology that occurred during research for a completely different radiation injury, there is a lot of uncertainty in the results due to the lack of a well-defined experimental design. Many secondary factors such as diet were not controlled and need to be. Further study could be done by minimizing explanatory variables and statistically analyzing their significance on the response variable – stomach distention. Also, a more quantifiable analysis could be performed if stomach volume could be recorded along with the explanatory variables over time. From this data, a better model could be formed to further benefit anyone performing a pre-clinical study with a similar methodology to this study.

For those performing similar studies, it is highly recommended to create shielding with dimensions that tightly contour the borders of the thorax (i.e. the lungs). We continued to use the 7.5 mm x 15 mm shield aperture since it guaranteed to expose a large portion of the lung without exposing areas beyond the thorax. If possible, a mold would be beneficial to immobilize the mice and minimize uncertainty in positioning; however, time and resources prevented us from obtaining such a mold.

## Figure Captions

*Figure 6*: Histology slides with Hematoxylin and Eosin staining at 20x multiplication. [A] A control mouse’s glandular stomach showing the three main layers: basal layer, middle layer, and superficial layer. [B] Similar stomach slice location for 15 week post-irradiation mouse. The middle layer’s parietal cells, which secretes acidic fluids, appears necrotic and poorly structured. An apparent ulcer appears in the basal layer of the stomach wall (green arrow). This indicates the desquamation of mucosa that protected the stomach wall from acid erosion and injury to the mucosa-secreting cells of the foveolar column.

